# Days Gained Response Discriminates Treatment Response in Patients with Recurrent Glioblastoma Receiving Bevacizumab-based Therapies

**DOI:** 10.1101/752402

**Authors:** Kyle W. Singleton, Alyx B. Porter, Leland S. Hu, Sandra K Johnston, Kamila M. Bond, Cassandra R. Rickertsen, Gustavo De Leon, Scott A. Whitmire, Kamala R. Clark-Swanson, Maciej M. Mrugala, Kristin R. Swanson

**Author notes:** Funding: Ben and Catherine Ivy Foundation, James T. McDonnell Foundation Grant (220020400TT), NIH Grants U54 CA210180, U54 CA143970, U01 CA220378, and R01 NS060752. Corresponding Author: Kyle W. Singleton, 5777 East Mayo Boulevard, Phoenix, AZ 85054.

## Abstract

**Purpose:** Accurate assessments of patient response to therapy are a critical component of personalized medicine. In glioblastoma multiforme (GBM), the most aggressive form of brain cancer, tumor growth dynamics are heterogenous across patients, complicating assessment of treatment response. This study aimed to analyze Days Gained (DG), a burgeoning model-based dynamic metric, for response assessment in patients with recurrent GBM who received bevacizumab-based therapies.

**Experimental Design:** Days Gained response scores were calculated using volumetric tumor segmentations for patients receiving bevacizumab with and without concurrent cytotoxic therapy (N=62). Kaplan-Meier and Cox proportional hazards analyses were implemented to examine DG prognostic relationship to overall (OS) and progression-free survival (PFS) from the onset of treatment for recurrent GBM.

**Results:** In patients receiving concurrent bevacizumab and cytotoxic therapy, Kaplan-Meier analysis showed significant differences in OS and PFS at previously identified DG cutoffs consistent with previous DG analyses using gadolinium-enhanced T1 weighted MR imaging. DG scores for bevacizumab monotherapy only approached significance for PFS. Cox regression showed that increases of 25 DG were significantly associated with a 12.5% reduction in OS hazard for concurrent therapy patients and a 4.4% reduction in PFS hazard for bevacizumab monotherapy.

**Conclusion:** Days Gained has significant meaning in recurrent therapy as a metric of treatment response, even in the context of anti-angiogenic therapies. This provides further evidence supporting the use of DG as an adjunct response metric that quantitatively connects treatment response and clinical outcomes.

## Introduction

In the era of precision-based medicine, clinicians strive to understand the unique evolution of disease in individual patients in order to provide the most effective care. Evaluation of treatment response and outcomes is of particular importance in clinical trials for comparing the effectiveness of novel therapies to the current standard of care. In the clinical setting, these same measurements of treatment response would ideally help clinicians assess tumor status in a timely manner and make informed decisions about adjusting therapies. In patients with glioblastoma multiforme (GBM), the most common and aggressive form of glioma, assessing tumor response to therapy has proven difficult. In recent years, the Response Assessment in Neuro-Oncology (RANO) working group has provided useful criteria for standardizing the assessment of response of high-grade gliomas to treatment, but there continues to be a discussion on how to expand on these guidelines by considering data from advanced imaging, digital subtraction maps, and volumetric measurements (1,2). Despite aggressive therapy, the highly invasive and dynamic nature of GBM inevitably leads to tumor recurrence, usually as defined by the existing response metrics, at which time previously useful therapies are rendered ineffective making determinations of response to subsequent therapies difficult. To address the challenge of treatment appraisal in the setting of recurrent GBM, we evaluated patients receiving bevacizumab with and without concurrent cytotoxic therapies using a personalized model-based response metric, Days Gained, that utilizes volumetric image measurements to account for differing tumor growth dynamics between patients.

## Background

Glioblastoma is the most common and aggressive infiltrating primary brain tumor. Patients’ typical length of survival from the time of diagnosis is less than two years (3,4). For patients 70 years of age and older, length of survival declines significantly to less than one year after diagnosis (5). Standard-of-care treatment consists of oral chemotherapy with alkylating agents and concomitant radiation therapy for a total of six weeks; adjuvant chemotherapy is then recommended for 6 to 12 months in the absence of disease progression or toxic side effects (6). To date, the management of recurrent GBM has posed significant challenges with limited success in clinical trials for the last several decades.

Bevacizumab, a humanized monoclonal antibody that targets vascular endothelial growth factor (VEGF), has been used to treat a number of cancers by inhibiting angiogenesis, thereby reducing the tumor’s innately dense and disorganized vascular supply. Bevacizumab is often used in combination with other cytotoxic therapies, such as irinotecan (topoisomerase 1 inhibitor) in colorectal carcinoma and paclitaxel (microtubule stabilizing agent) in breast cancer. Early reports from the AV37018g and NCI 06-C-0046E trials of bevacizumab for recurrent GBM were promising for improved progression-free survival (PFS), but improvements were not observed for overall survival (OS) (7,8). These trials resulted in FDA approval of bevacizumab as a single-agent treatment for recurrent GBM. While subsequent clinical trials have not provided conclusive evidence that bevacizumab improves OS, they have solidified the impact of bevacizumab on PFS and clinicians continue to use it both as a monotherapy and in combination with cytotoxic agents (9).

Primary measurements of treatment response from the Macdonald and RANO criteria use two-dimensional tumor measurements to assess changes in contrast enhancement on T1-weighted gadolinium enhanced (T1Gd) magnetic resonance imaging (MRI) (1,2,10). As bevacizumab inhibits neoangiogenesis and normalizes the blood-brain barrier within the tumor, gadolinium extravasation is diminished and contrast enhancement on imaging diminishes. These effects can be visualized as early as 1-2 days after therapy and can persist for the duration of bevacizumab administration. As such, assessing the efficacy of bevacizumab with imaging has proven difficult due to this “pseudo-response” effect, where imaging response may reflect anti-angiogenic response rather than significant cytoreduction in tumor cell burden. To combat this shortcoming in response assessment that originated during use of the Macdonald Criteria, RANO incorporated additional guidelines that considered non-enhancing tumor progression visualized on other edema-capturing imaging sequences, such as T2-weighted (T2) and T2 fluid attenuated inversion recovery (FLAIR) sequences on MRI. In one study, it was noted that as many as 37% of patients receiving bevacizumab had tumor recurrence defined specifically by T2/FLAIR changes (11). Under the current paradigm, patients are considered to have failed treatment if their T1Gd enhancing imaging abnormality has increased by 25% or more. However, determining changes in the size of a non-enhancing (T2/FLAIR) abnormality remains a subjective assessment. The RANO working group considers T2/FLAIR assessment to be a major challenge for the field due to the difficulty in measuring non-enhancing regions accurately (2). Thus, exploration of additional response metrics is warranted in order to address these complex challenges.

Clinicians have few effective tools for assessing treatment response in a clinically-relevant timeframe that are also consistently predictive of outcomes for the patient (1,12). Current imaging-based treatment response metrics in cancer utilize one-dimensional (RECIST) or two-dimensional (Macdonald, RANO) measurements of tumor abnormality (1,2,10,13,14). These measurements capture only a portion of the total abnormality seen on MRI and do not represent the entire scope of disease for each patient. As a result, current metrics are limited in their ability to describe patient-specific differences in tumor size and growth in GBM and have shown little success in predicting patient outcome (1). The kinetics of tumor growth have been shown to vary greatly across patients due to the heterogeneous nature of GBM. Consequently, developing response metrics that account for tumor kinetics can aid in the understanding of tumor aggressiveness that have not been taken into account with current metrics (15,16).

A number of mathematical models have been previously investigated for the purpose of simulating tumor growth kinetics (17–22). These mathematical models integrate volumetric clinical imaging data to generate patient-specific simulations that forecast cell densities in the tumor environment. Such simulations, therefore, provide untreated virtual controls (UVCs) for each patient that can be used to estimate anticipated growth at future timepoints for comparison against actual tumor growth on post-treatment imaging. Utilizing this model-based approach, a patient-specific metric, Days Gained (DG), was defined as the degree to which a given treatment delayed tumor growth, measured in days (23). DG was initially applied in the context of first line, standard-of-care radiotherapy and was found to be prognostic for both OS and PFS (23). Using additional cases in the first-line radiotherapy setting, DG was further assessed for sensitivity to the complexity of the UVC tumor model (24). Three models with different levels of computational complexity (four-dimensional anatomic, four-dimensional spherical, and linear) were used to compute Days Gained scores. In each case, DG remained prognostic for OS and PFS, indicating that simplified versions of the UVC were appropriate for use in response metric assessment. Simplifying the UVC model can greatly reduce computational time and makes a model-based response metric like DG more readily accessible to clinicians. In addition, this work found that DG was able to discriminate progression versus pseudo-progression following radiotherapy. Following these works, DG was also applied in a novel early phase gene therapy clinical trial using autologous gene-modified hematopoietic stem cells (25). DG scores were calculated for patients receiving therapy and for associated controls receiving standard of care therapy. DG values were higher for patients undergoing the novel therapy indicating this treatment caused a greater deflection of tumor growth than standard care.

Based upon these prior successes, we attempt to further elucidate the capability of DG in the recurrent setting where treatments begin to vary and can be given in combination. The incorporation of patient-specific kinetics into metrics of response can allow for the separation of prognostically-predictive treatment effects from GBM heterogeneity. As noted above, bevacizumab use for the treatment of GBM has been widespread since FDA approval, but there remains no conclusive data that bevacizumab alone can improve overall survival. Consequently, bevacizumab is frequently given with other therapies, but the pseudo-response effect of bevacizumab can impair assessment of response. Thus, we investigate a cohort of patients who received bevacizumab as monotherapy or in combination with cytotoxic therapies using the Days Gained response metric to evaluate discrimination of OS and PFS outcomes.

## Methods

Following institutional review board approval, we identified 67 patients diagnosed with recurrent glioblastoma who received bevacizumab therapy with or without concurrent cytotoxic therapy from our multi-institutional clinical research database. In addition, inclusion criteria required each patient to have T1-weighted gadolinium enhanced (T1Gd) and T2 fluid attenuated inversion recovery (FLAIR) magnetic resonance images on two pre-treatment and one post-treatment date from the start of bevacizumab therapy. We reviewed the treatments of each recurrent GBM patient to determine what concurrent cytotoxic treatment, if any, were given alongside bevacizumab therapy. Patients in our dataset received concurrent carboplatin, CCNU (lomustine), BCNU (carmustine), Gliadel Wafers, or CPT-11 (irinotecan) with bevacizumab. Carboplatin was the most common concurrent cytotoxic therapy administered (N=25). Patients were analyzed as a complete set of bevacizumab-based therapies and were also analyzed as subgroups of bevacizumab monotherapy (BevAlone, N=24) and bevacizumab plus cytotoxic agent (BevCyto, N=38).

Patient response to therapy was evaluated using our previously-described Days Gained response metric (23,24). Briefly, Days Gained is computed by comparing post-treatment volumetric tumor size against a patient-specific prediction of untreated tumor size over the period of time between imaging studies. Patient-specific predictions are obtained using tumor growth characteristics calculated using spherically equivalent radial tumor size as measured from the volumetric segmentations of tumor abnormality. Volumetric segmentations were performed for T1Gd and FLAIR images in our cohort using a semi-automated segmentation process by one or more independent image analysts trained in MRI segmentation. Days Gained scores were calculated using the volumetric growth characteristics measured from each imaging sequence type.

For each treatment group and imaging sequence type, we performed Kaplan-Meier analysis as well as univariate and multivariate Cox proportional hazards models for progression free survival (PFS) and overall survival (OS). Kaplan-Meier analysis was first performed on the full dataset using previous cutoffs from DG evaluation in patients receiving radiotherapy (24). Previous optimal cutoffs were then applied in the treatment subgroups and were also compared to median DG cutoffs. Univariate Cox proportional hazards was performed for DG and multivariate Cox analysis for DG, age at start of therapy, and patient sex. Kaplan-Meier and Cox proportional hazards analysis were performed in R (v3.6.1) using the *survival* and *survminer* packages (26,27). OS and PFS were defined as the time interval between the start of the patient’s recurrent therapy to the date of death or date of progression, respectively, as documented in the clinical chart. Patients were censored at last known date of follow-up if the outcome in question was not available.

## Results

### Patient Cohort

A total of 62 patients with recurrent GBM met criteria for inclusion in this study. Five cases were removed due to a lack of required imaging timepoints. Of these patients, 24 received bevacizumab monotherapy (BevAlone) and 38 received bevacizumab with concurrent cytotoxic therapy (BevCyto). Days Gained scores were calculable from T1Gd imaging (DG_T1Gd_) for 62 patients and for 53 patients from FLAIR imaging (DG_FLAIR_). A summary of patient demographics and calculated Days Gained scores for T1Gd and FLAIR imaging are provided in Table 1.

**Table 1:**
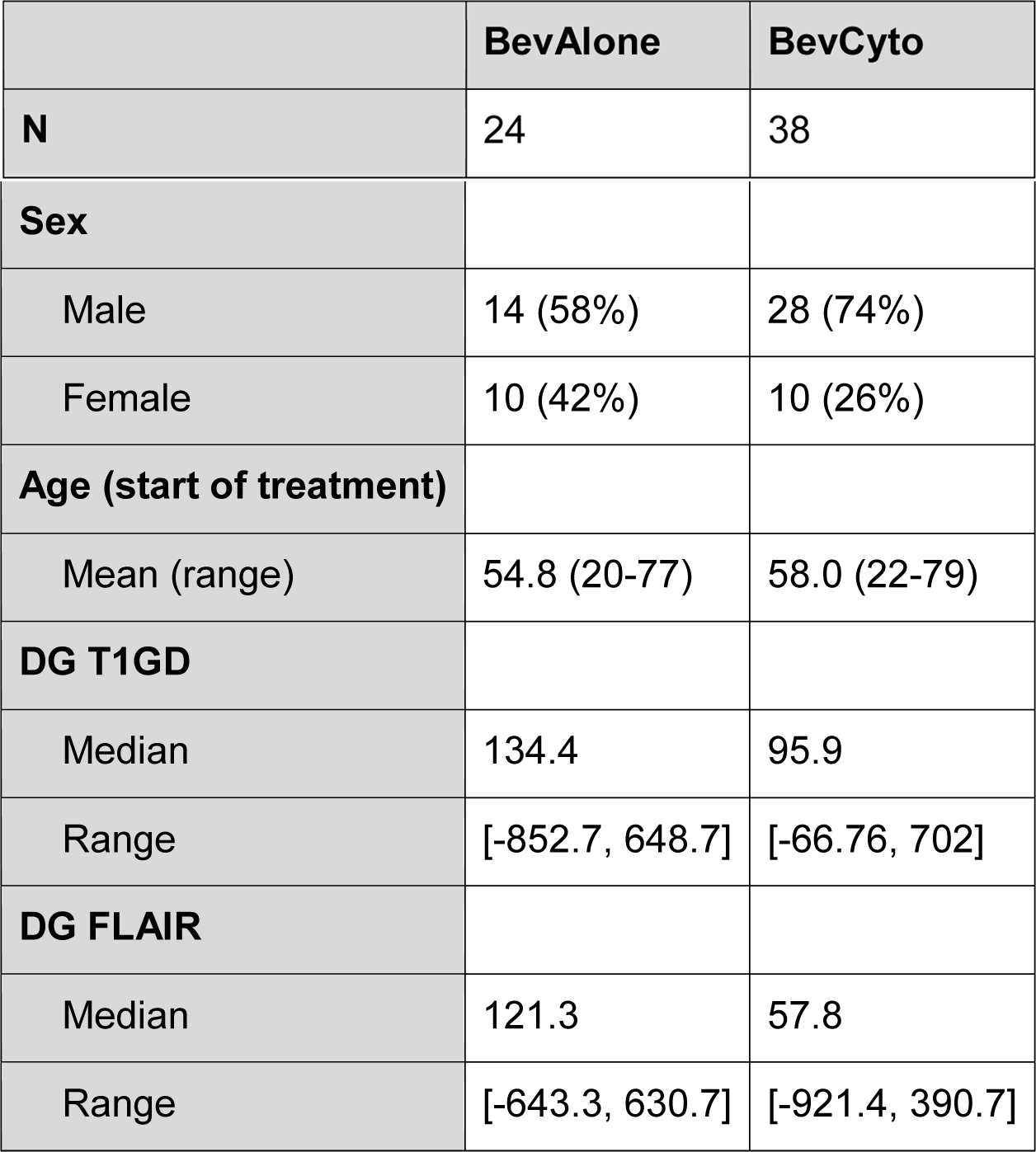
Demographics of patients with recurrent glioblastoma by treatment group evaluated with Days Gained scores. Patients received either bevacizumab monotherapy (BevAlone) or bevacizumab concurrent with a cytotoxic therapy (BevCyto).

### Prior DG Cutoffs Remain Significant in Bevacizumab treated Recurrent Patients

In prior analysis of newly diagnosed GBMs receiving standard-of-care therapies, a range of DG scores (OS: [65, 105] and PFS: [55, 110]) were identified with statistically significant survival benefit (23,24). Reported optimal thresholds in these ranges (78 DG for OS and 93 DG for PFS using T1Gd imaging) were applied to the overall bevacizumab-treated cohort of 62 recurrent GBM patients. Each cutoff continued to successfully distinguish OS and PFS using DG_T1Gd_ scores (Figure 1, log-rank p-values < 0.0001). Cutoffs were not significant for DG_FLAIR_ scores (Supplement SF1).

**Figure 1:**
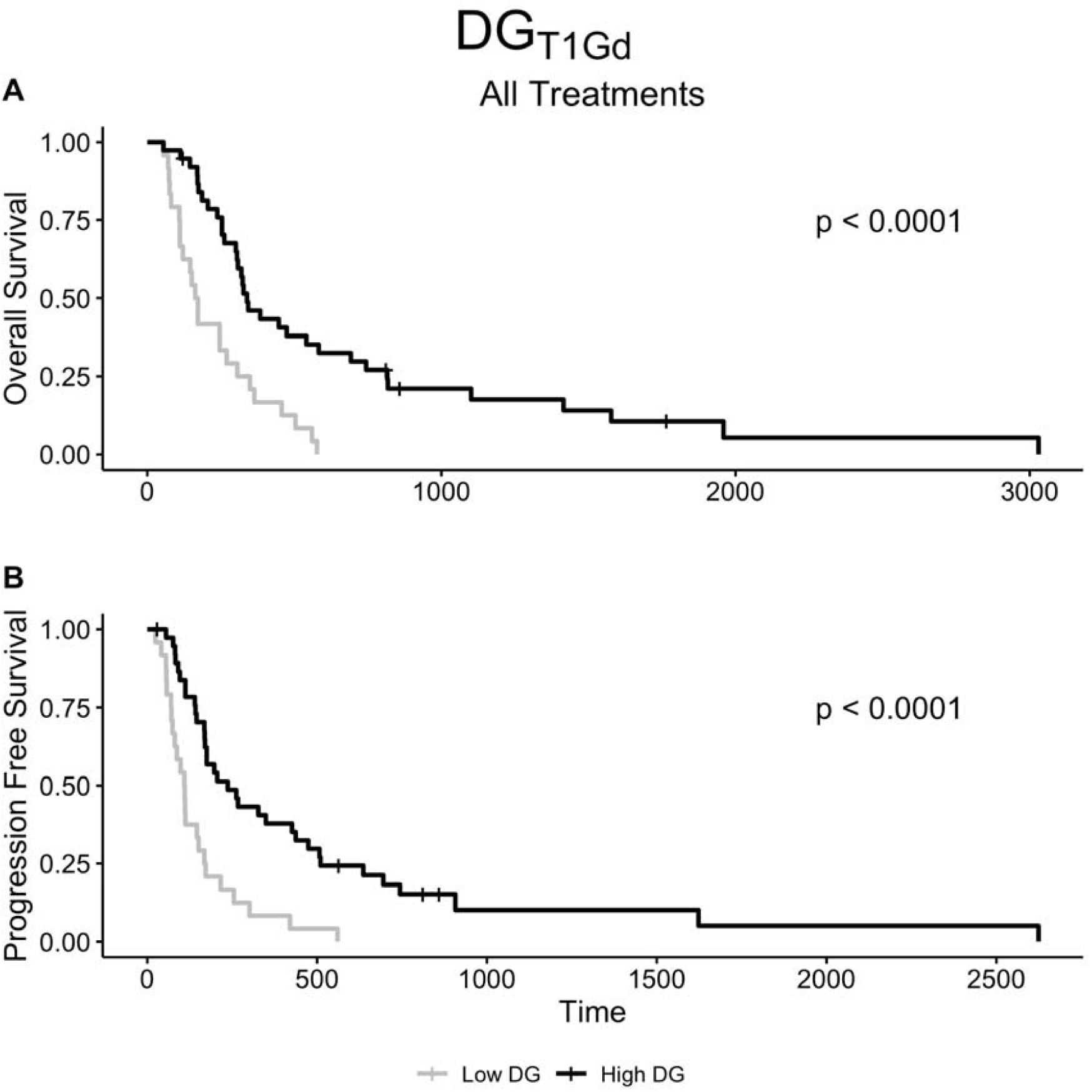
Kaplan-Meier analysis applying previously identified optimal DG_T1Gd_ thresholds from upfront GBM patients (78 DG, OS; 93 DG, PFS) to recurrent bevacizumab-treated cohort with DG_T1Gd_ scores. Prior thresholds significantly discriminated survivor groups using DG_T1Gd_ in the recurrent setting.

### Subanalysis of Patients treated with Cytotoxic Therapies in Combination with Bevacizumab

To further consider a generalized target DG cutoff, prior DG thresholds were compared with a median dichotomization in a subanalysis of treatment groups. A comparison of the prior and median cutoffs is provided in Table 2. Median DG_T1Gd_ scores for BevCyto patients were highly significant for both OS (p=0.00036) and PFS (p=0.00085) in Kaplan-Meier analysis log-rank tests (Table 2, Figure 2B and 2D). Prior thresholds were also highly significant for OS (p<0.0001) and PFS (p=0.00039). In contrast, the BevAlone group was not significant (OS, p=0.182; PFS, p=0.059) in Kaplan-Meier analysis using either the median or prior cutoffs (Table 2, Figure 2A and 2C). In addition, when using DG_FLAIR_ scores to assess response, the BevAlone or BevCyto groups did not show a significant difference between the high and low DG cutoff for either threshold (Supplement SF2).

**Table 2:**
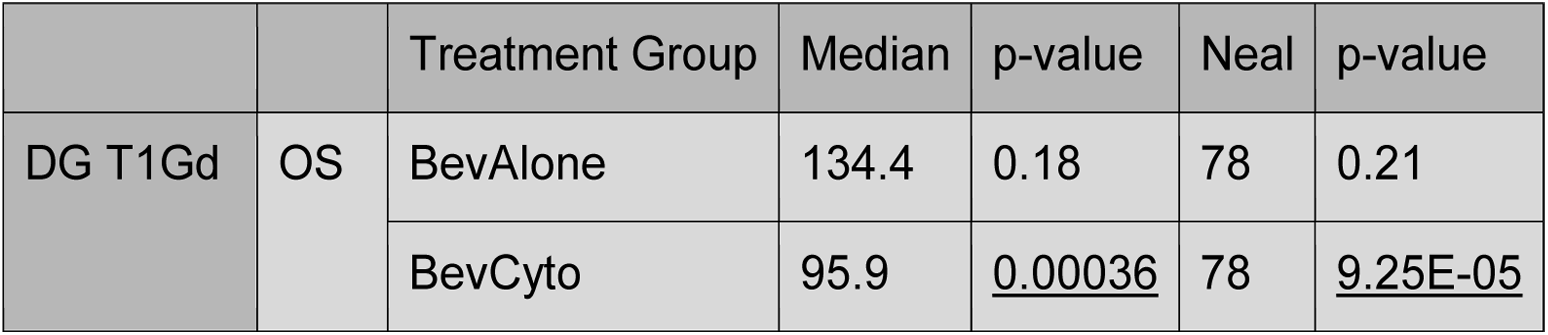
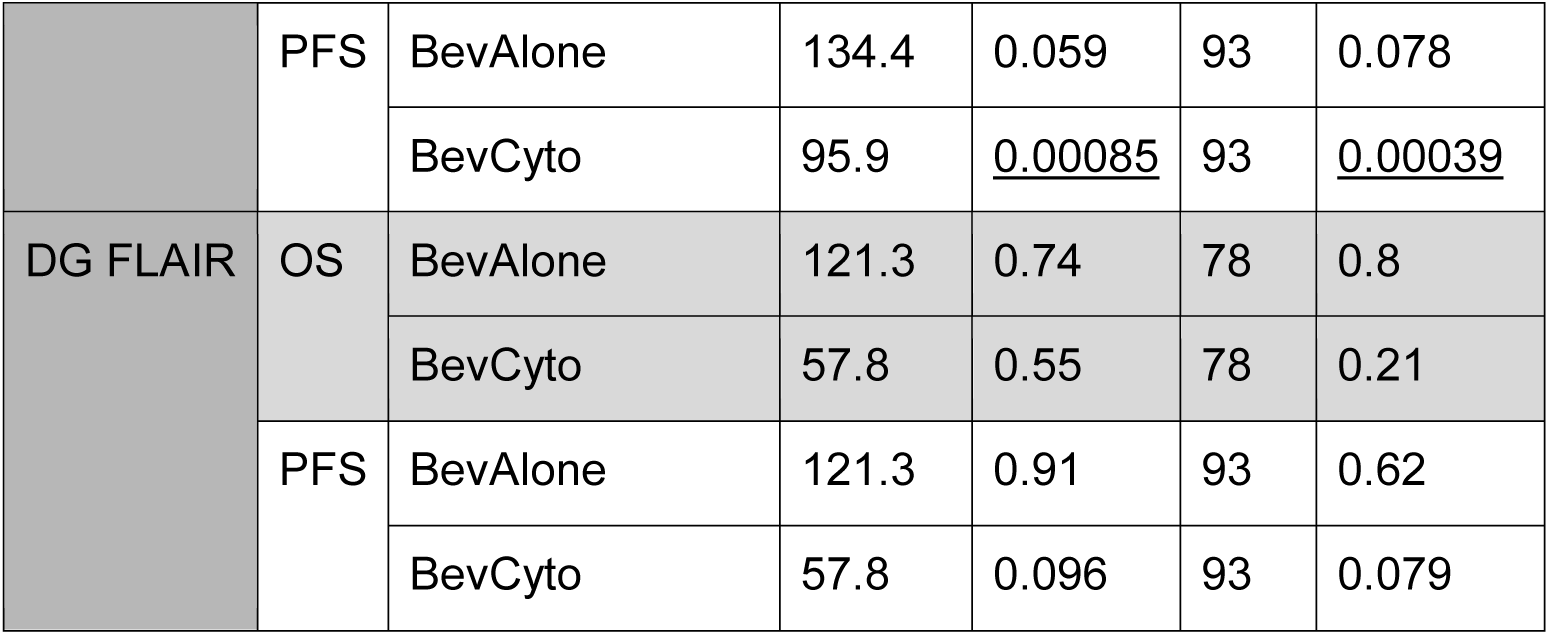
Efficacy of median, optimal, and previously reported optimal thresholds (Neal: 78 DG OS, 93 DG PFS (24)) for discriminating OS and PFS across all treatment groups for both T1Gd and FLAIR based DG scores. Significant log-rank test p-values underlined.

**Figure 2:**
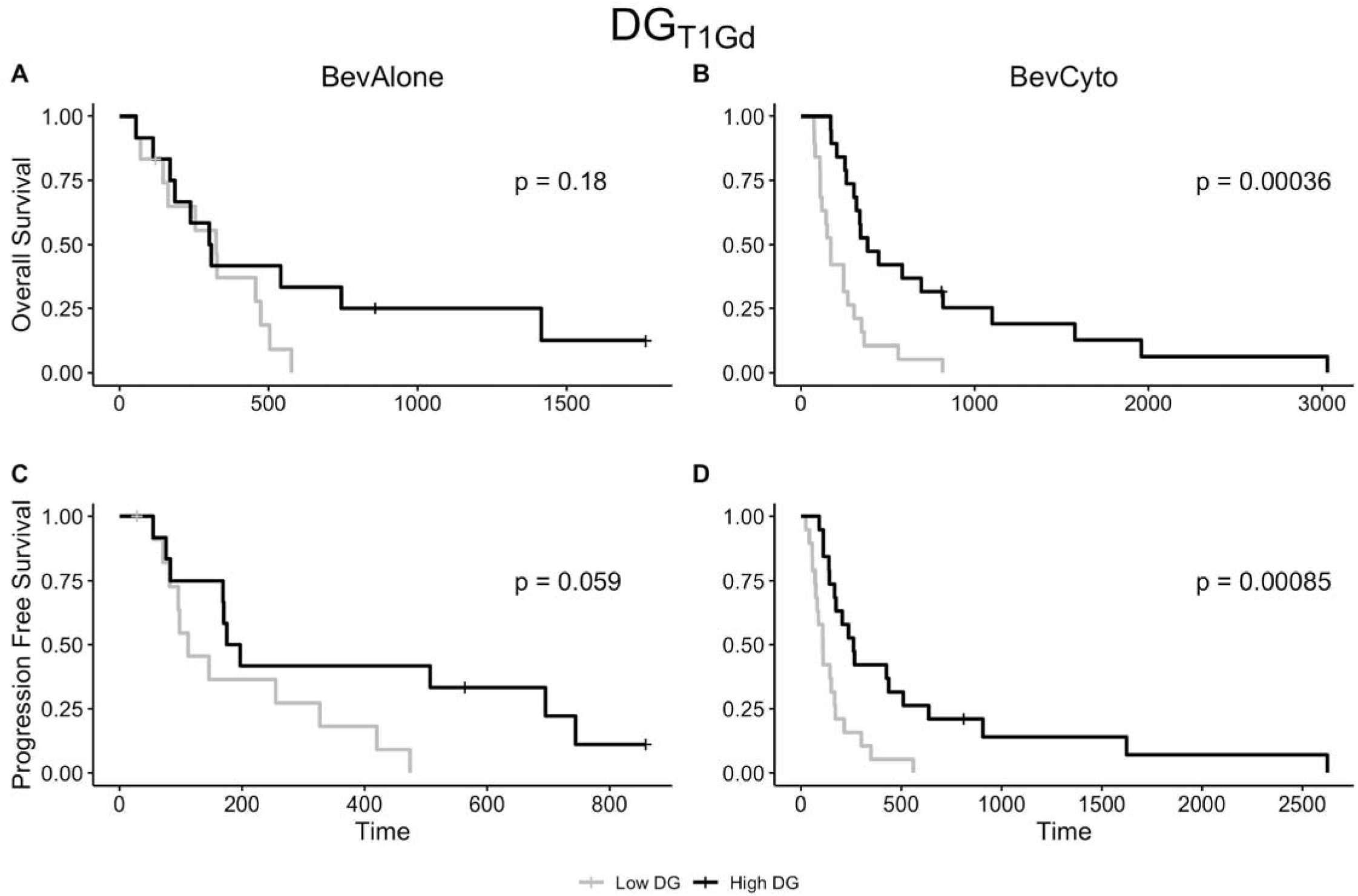
Median Kaplan-Meier plots of DG_T1Gd_ by treatment group for overall survival (top row, A-B) and progression free survival (bottom row, C-D). Dichotomizing the population at the median DG_T1Gd_ in the bevacizumab alone treatment group, failed to distinguish OS but was near significant for PFS. In the bevacizumab and concurrent cytotoxic therapy group, the median distinguished survival.

### Robustness of DG as a Predictor of Survival

Using an iterative Kaplan-Meier analysis, we further explored the range of Days Gained thresholds that discriminate for OS and PFS in each therapy group. Figure 3 illustrates these results for OS and PFS across both treatment groups using DG_T1Gd_ thresholds. DG_T1Gd_ thresholds that reached significance were broadly seen in a range between 20 to 250 DG_T1Gd_ for OS in BevCyto patients. This range was consistent for PFS. In addition, these significant DG ranges overlap with previous findings for Days Gained cutoffs in newly diagnosed patients receiving upfront therapy, where significant DG thresholds ranged from 65 to 105 for OS and 55 to 110 for PFS (Figure 3, red box) (24). In the BevAlone group, no significant DG_T1Gd_ thresholds were seen for overall survival for BevAlone patients. A few significant thresholds for BevAlone PFS were observed around 60 and 140 DG for T1Gd and FLAIR analysis, but other intermediate thresholds were not significant. Thresholds for DG_FLAIR_ analysis were not significant in most cases, although a few significant thresholds were observed for PFS in the BevCyto therapy group (Supplement SF3).

**Figure 3:**
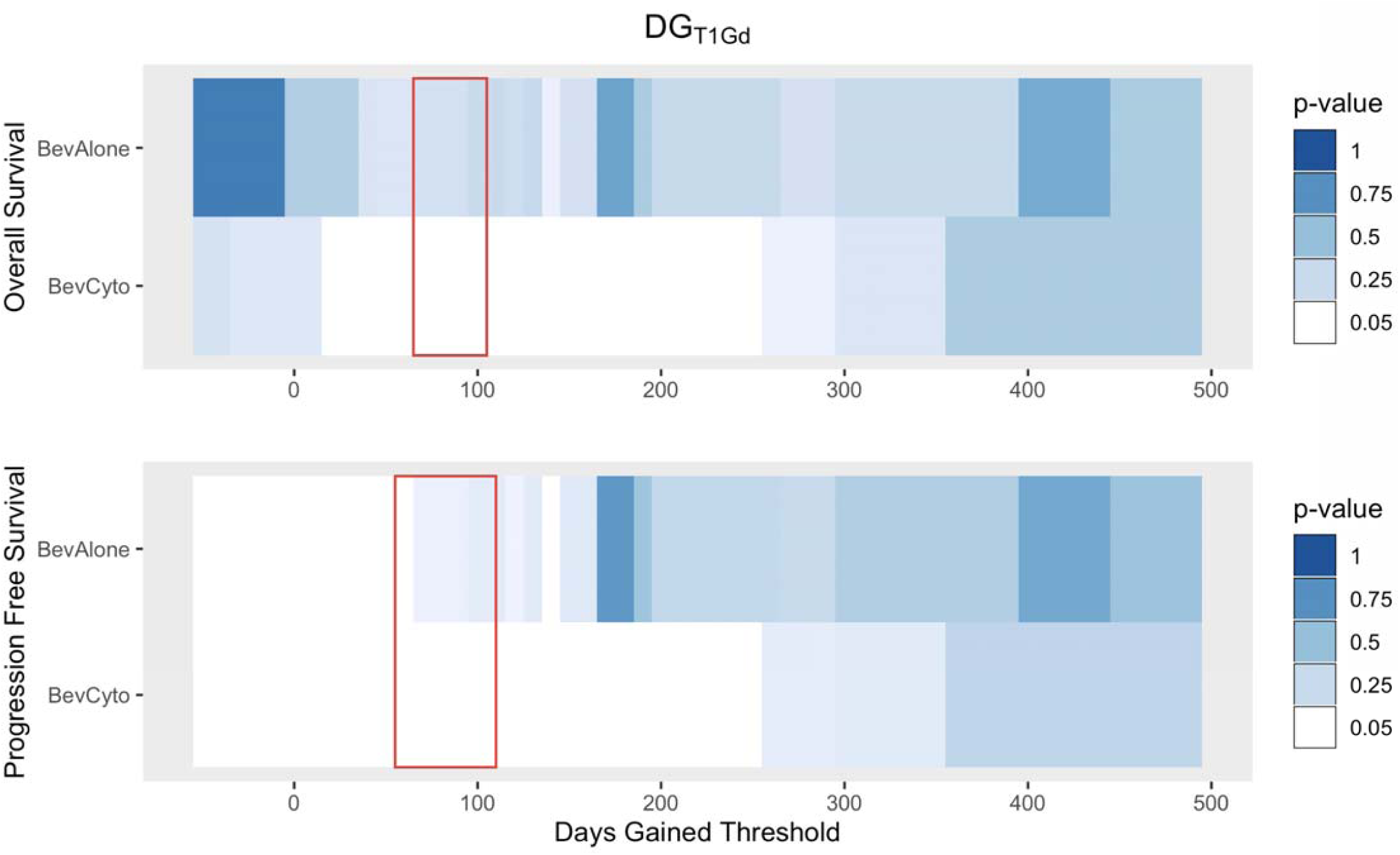
Iterative Kaplan-Meier significance of DG_T1Gd_ thresholds for overall survival and progression free survival for each therapy group (white: log-rank p<=0.05, blues: p>0.05). Cytotoxic therapy thresholds show substantial overlap with Days Gained thresholds from newly diagnosed treatment analysis (red box (24)).

### Cox Proportional Hazard Analysis of DG

In Cox proportional hazards regression analysis, DG_T1GD_ was a significant predictor of OS for the concurrent therapy groups in univariate analysis and after controlling for patient age and sex in multivariate analysis (Table 3). In the BevCyto group, for example, patients had a 12.5% reduction in the chance of death for every 25 Days Gained. Treatment with bevacizumab monotherapy was not a significant predictor of patient OS, but was significant for PFS (Table 3). Patients in the BevAlone gr up saw a 4.4% reduction in the chance of progression for every 25 DG. Similar trends were seen for PFS in the concurrent therapy group, with a reduction of 12.3% for BevCyto. Male sex was also a signficant predictor of decreased survival in some models (Table 3). DG_FLAIR_ scores were not significant as a predictor of patient OS or PFS in either treatment group (Supplement ST1).

**Table 3:**
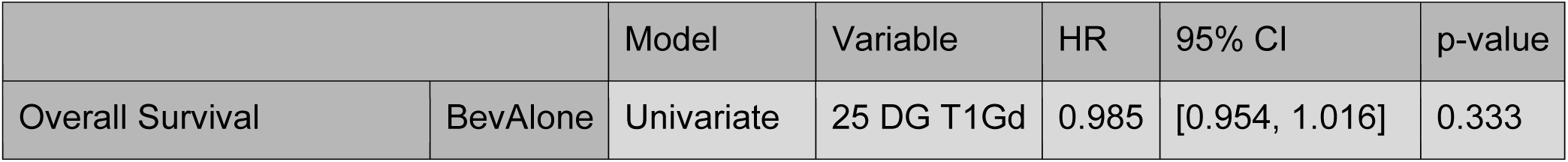
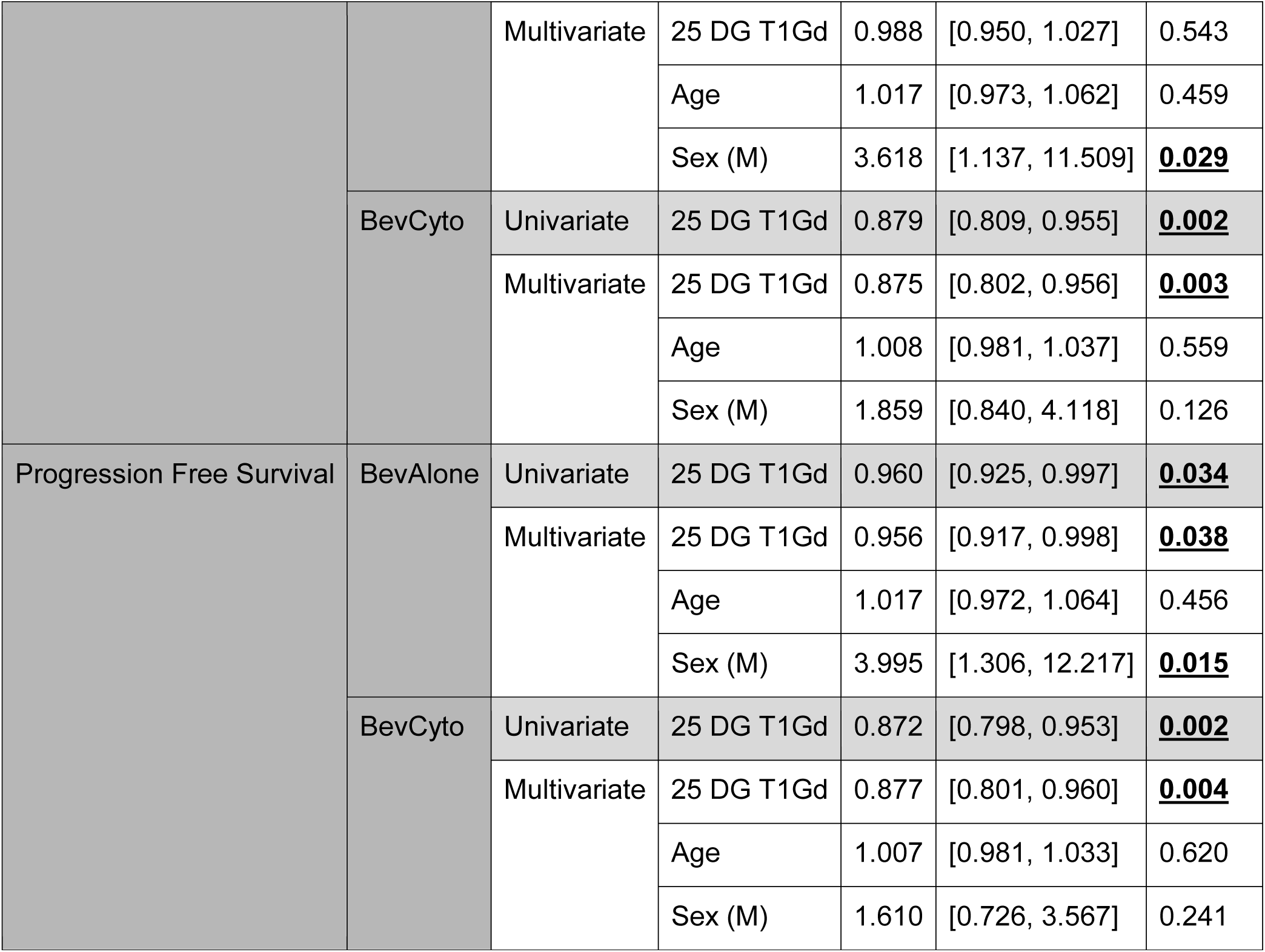
Cox proportional hazards regression analysis of overall survival and progression free survival using continuous DG_T1Gd_ scores, patient age at start of treatment, and patient sex. Significant p-values underlined.

## Discussion

In this work, we examined the use of a patient-specific response metric, Days Gained, for the discrimination of OS and PFS in patients with recurrent GBM who received bevacizumab with or without concurrent cytotoxic therapy. Patients were grouped by therapy to examine the possible pseudo-responsive effects from bevacizumab on response scoring using Days Gained. Patients receiving concurrent cytotoxic therapy with bevacizumab were most similar to prior DG analysis in patients receiving adjuvant radiotherapy (23,24). Days Gained scores in this group showed similar results to prior adjuvant findings and thresholds applied from the previous adjuvant analysis (24) remained significant in this recurrent bevacizumab-based setting for OS and PFS (Table 2, Figure 2). Significant OS and PFS benefit was also demonstrated for concurrent therapy patients in Cox regression analysis for increases in DG scores (Table 3). Patients receiving bevacizumab monotherapy showed a different pattern of Days Gained response with significant DG thresholds found only for PFS in both Kaplan-Meier (Table 2, Figure 2) and Cox proportional hazards regression analysis (Table 3). These results support Days Gained as a patient-specific metric of response that can be used to better inform clinical decision-making with regard to treatment course.

Two randomized clinical trials investigated the addition of bevacizumab to standard of care treatment, temozolomide and radiotherapy, in nearly 1100 patients with GBM (28,29). In both studies, the time to clinical progression was longer in patients who received concurrent bevacizumab than standard therapy. However, there were no differences in overall survival between groups. In light of these results, it is unsurprising that DG scores were unable to discriminate OS in patients receiving bevacizumab monotherapy in our study.

Days Gained has been used previously to discriminate OS and PFS in patients with newly diagnosed GBM who have received radiation as a part of standard-of-care treatment (23,24). Kaplan-Meier analysis from that work revealed a range of significant Days Gained thresholds. In our study, significant DG thresholds spanned overlapping ranges for OS and PFS in Kaplan-Meier analysis. More importantly, prior optimal thresholds from the adjuvant setting successfully discriminated OS and PFS in the full population of recurrent patients receiving bevacizumab-based therapy. The overlap of DG thresholds from this work with prior findings and applicability of past thresholds to these recurrent cases further emphasizes that DG scores provide a stable marker, useful for discrimination across different points of patient care, even in the challenging context of bevacizumab-based therapies that are known to significantly impact imaging changes but with unclear benefit in overall survival outcomes.

While a pooled analysis of DG for all cases from our BevAlone and BevCyto cohorts was significant with regard to discriminating outcomes, the majority of discrimination effect was related to patients receiving concurrent cytotoxic therapy with bevacizumab. Concurrent therapy patients could be discriminated into survival groups using DG_T1Gd_ scores with high significance, but bevacizumab monotherapy only denoted a detectable benefit for PFS (Table 2, Figure 2). For cases including concurrent bevacizumab therapy, these findings also indicate that treatment response is detectable using DG scores in the presence of potential pseudo-response effects. This stands in contrast to clinical trials that have found no benefit to combination bevacizumab therapy. For example, a study of progressive GBM compared lomustine, a common therapy at first progression, with lomustine in combination with bevacizumab (30). The addition of bevacizumab conferred a prolongation of PFS without an overall survival advantage. Our findings indicate there may be subpopulations of patients who benefit from combination therapy that are difficult to detect. Days Gained appears to be able to help detect these individual patients, but this requires further study.

Cox proportional hazards modeling further supports Days Gained as a significant indicator of OS and PFS in assessing the utility of bevacizumab in recurrent GBM and treatment response more broadly. Patients receiving concurrent therapy in this work saw a significant reduction in hazard of 12.5% per 25 DG_T1Gd_ for overall survival (Table 3). A similar reduction in hazard was seen for increasing Days Gained in PFS. A more modest 4.4% hazard reduction per 25 DG_T1Gd_ for PFS was also seen for the bevacizumab monotherapy group (Table 3). These findings demonstrate that increasing DG values relate to greater total benefit in outcomes from therapy, suggesting that tumor size reduction relative to tumor growth rate has significant meaning for predicting outcomes across the variety of anti-angiogenic and cytotoxic therapies administered. While glioblastoma ultimately remains fatal in all patients, being able to prognosticate based on the magnitude of a patient’s tumor specific kinetic properties provides clinical value. Our method may serve as a useful adjunct to the Macdonald and RANO criteria which is used to classify responders from non-responders to therapy. Days Gained offers to incorporate tumor individuality with regard to the dynamics of growth prior to treatment. Thus, assessing patients with Days Gained can provide additional context to overall tumor size and new opportunities to adjust followup times for each patient, particularly those with low Days Gained where response appears to be limited. Translating treatment and followup schedules based upon patient specific context may help clinicians gain traction towards the goal of connecting response to clinical benefit, as measured in terms of outcomes like overall survival.

Notably, DG calculated from FLAIR images did not show a strong relationship with OS or PFS for any treatment group in our analysis. Measuring T2/FLAIR abnormality can be difficult and has been noted by the RANO working group as a reason why objective criteria have not been added for non-enhancing image assessments in the RANO criteria (2). Pre-treatment growth measurements of FLAIR were more varied in our cohort than for T1Gd imaging due to the difficulty of measuring these images. As a result, this variability may have played a role in our analysis. Other studies have found a difference in OS and PFS between patients with differing radiologic progression patterns (11,31), suggesting additional analysis with T2/FLAIR imaging is likely warranted. In addition, as with assessment with the RANO criteria, assessment at future imaging timepoints after more cycles of bevacizumab therapy may be required to detect these differences between patients. We hope to add more cases and timepoints to future analysis using DG.

We acknowledge the innate limitations of performing a retrospective analysis of patients. Treatment options for recurrent GBM have varied significantly over time with attempts to improve patient outcomes, yet there is no defined standard-of-care for this recurrent context. We combined patients who received a variety of different concurrent treatments to serve as a cytotoxic treatment group, but the signal may be confounded by additional surgery or lack of efficacy of some treatments. For example, systemic nitrosoureas and other alkylating agents, such as lomustine, procarbazine, and vincristine, have been trialed in different combinations with no clear treatment benefit for the average patient in cohort studies (32–36). Patients receiving these treatments with concurrent bevacizumab may experience imaging changes similar to bevacizumab monotherapy cases, modulating apparent DG response in the group. However, our findings indicate that these effects do not limit DG ability to discriminate responders. Another limitation in our work is sample size. Given the heterogeneity in treatment profiles of recurrent patients and frequently non-standardized, varying intervals of scheduled MR imaging, we were limited in the number of cases we were able to include in our analysis. Prospective data collection and randomized clinical trials provide the greatest opportunity to validate our predictive findings with a larger sample size. Our current analysis provides evidence to support these future validations. In fact, only recently has the RANO criteria been assessed more formally using outcome data from trials (31). As a result, sample size remains a constant challenge for response evaluation. Nevertheless, advances in medical imaging and electronic health records have reduced the difficulty of obtaining required imaging and treatment information, which will only make application of DG and other personalized metrics easier in the future.

## Conclusion

Our study indicates that Days Gained, an individualized metric of response to therapy, was able to discriminate OS and PFS for patients receiving cytotoxic therapies in combination with bevacizumab, but could only discriminate PFS for those receiving bevacizumab alone. This finding is consistent with the growing literature that bevacizumab monotherapy does not impact OS. The discriminative power of DG for cytotoxic therapy using T1Gd MRI does not appear to be negatively affected by the administration of concurrent bevacizumab. Significant DG thresholds were consistent with prior DG findings and thresholds from these prior studies of standard-of-care therapy continued to significantly discriminate OS and PFS for recurrent cases. Overall, including in multivariate Cox proportional hazards regression analysis, increased DG scores were significantly associated with decreased progression in patients receiving bevacizumab-based therapies and increased survival in patients receiving bevacizumab concurrent with cytotoxic therapy. This evidence further supports DG as a clinically meaningful metric of individualized treatment response, even in the context of anti-angiogenic therapies that are known to ambiguously modulate imaging features on MRI.

## Acknowledgements

We would like to thank Destiney Kirby for contributions to the collation of cases for the dataset used in this work.

## Supplemental Figures and Tables

**Figure SF1:**
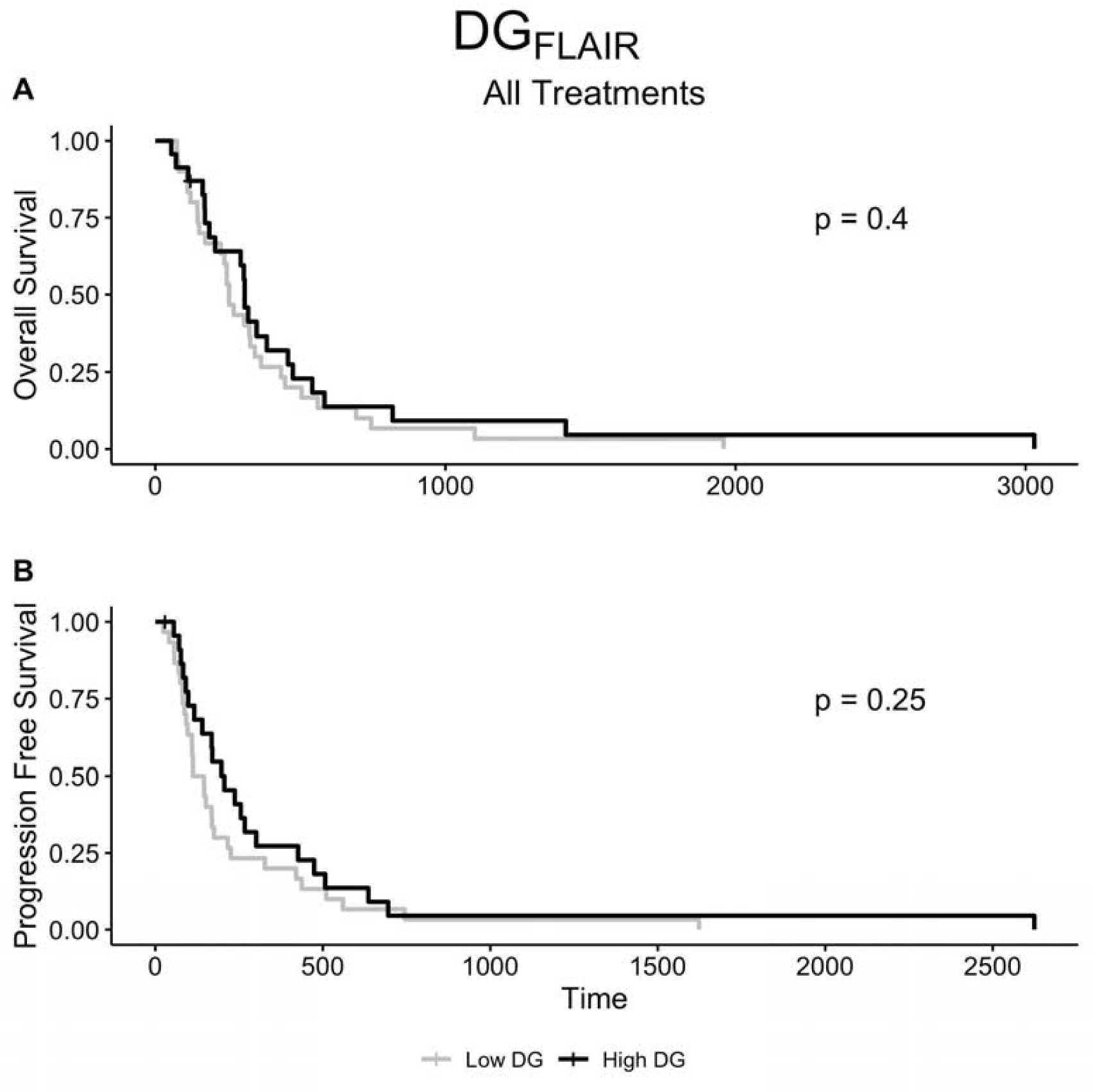
Kaplan-Meier analysis applying previously identified optimal DG_T1Gd_ thresholds from upfront GBM patients (78 DG, OS; 93 DG, PFS) to recurrent bevacizumab-treated cohort with DG_FLAIR_ scores. No significant discrimination between groups was observed.

**Figure SF2:**
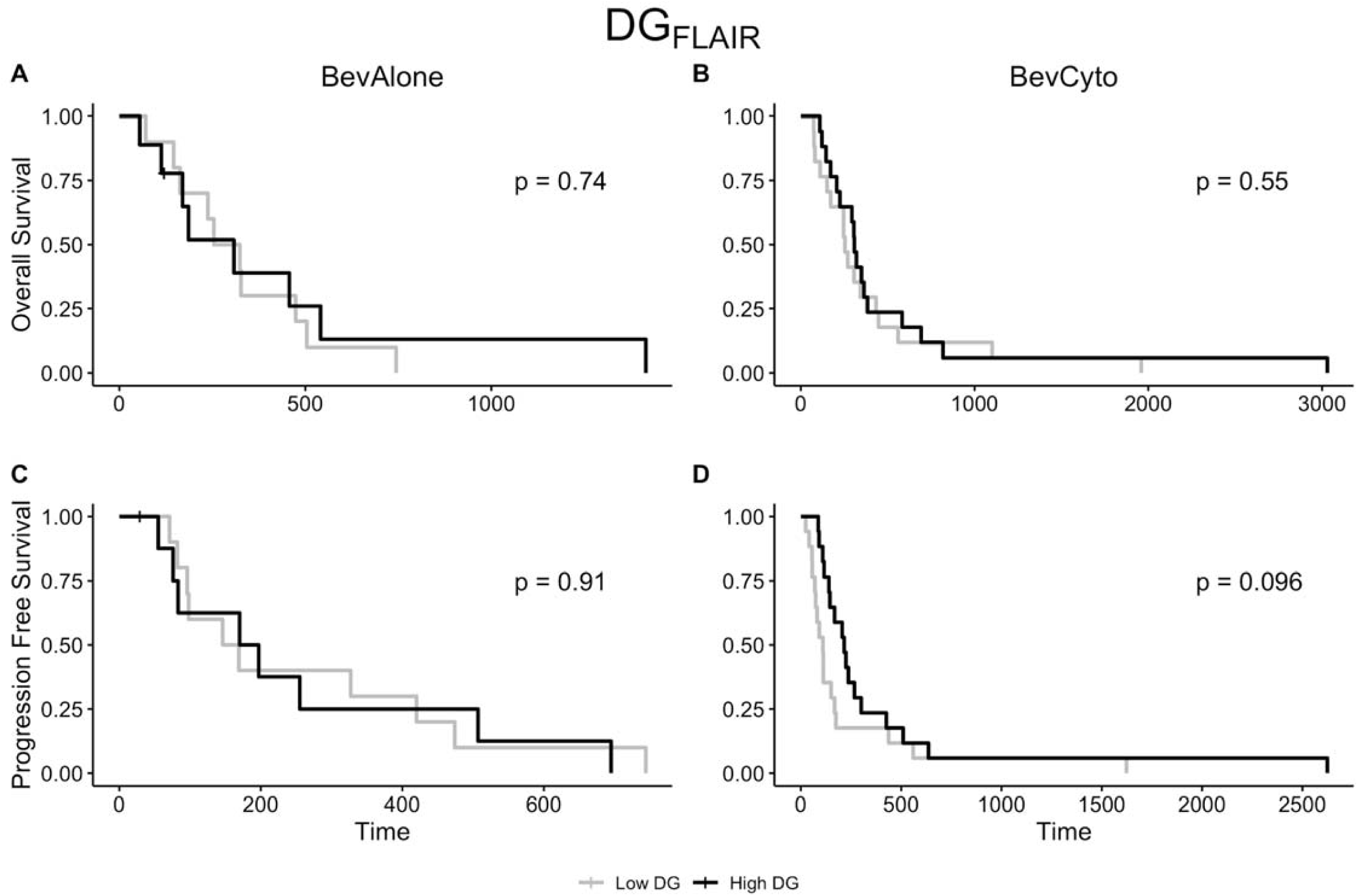
Median Kaplan-Meier plots of DG_FLAIR_ by treatment group for overall survival (top row, A-B) and progression free survival (bottom row, C-D). No significant discrimination between groups was observed.

**Figure SF3:**
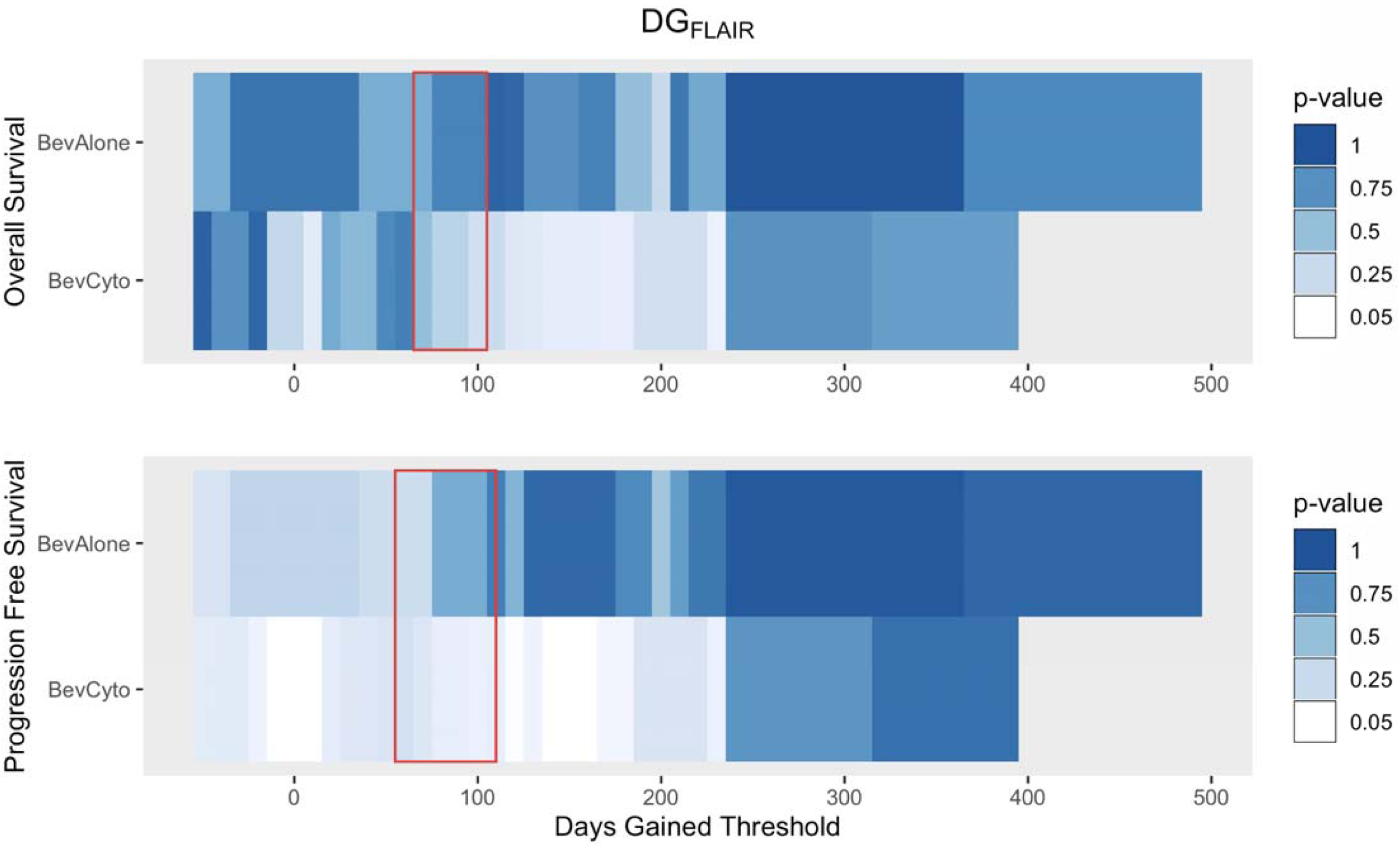
Iterative Kaplan-Meier significance of DG_FLAIR_ thresholds for overall survival and progression free survival for each therapy group (white: p<=0.05, blues: p>0.05). Very few thresholds overlap with Days Gained thresholds from prior adjuvant treatment analysis (red box (24)) based on T1Gd imaging.

**Table SF1:**
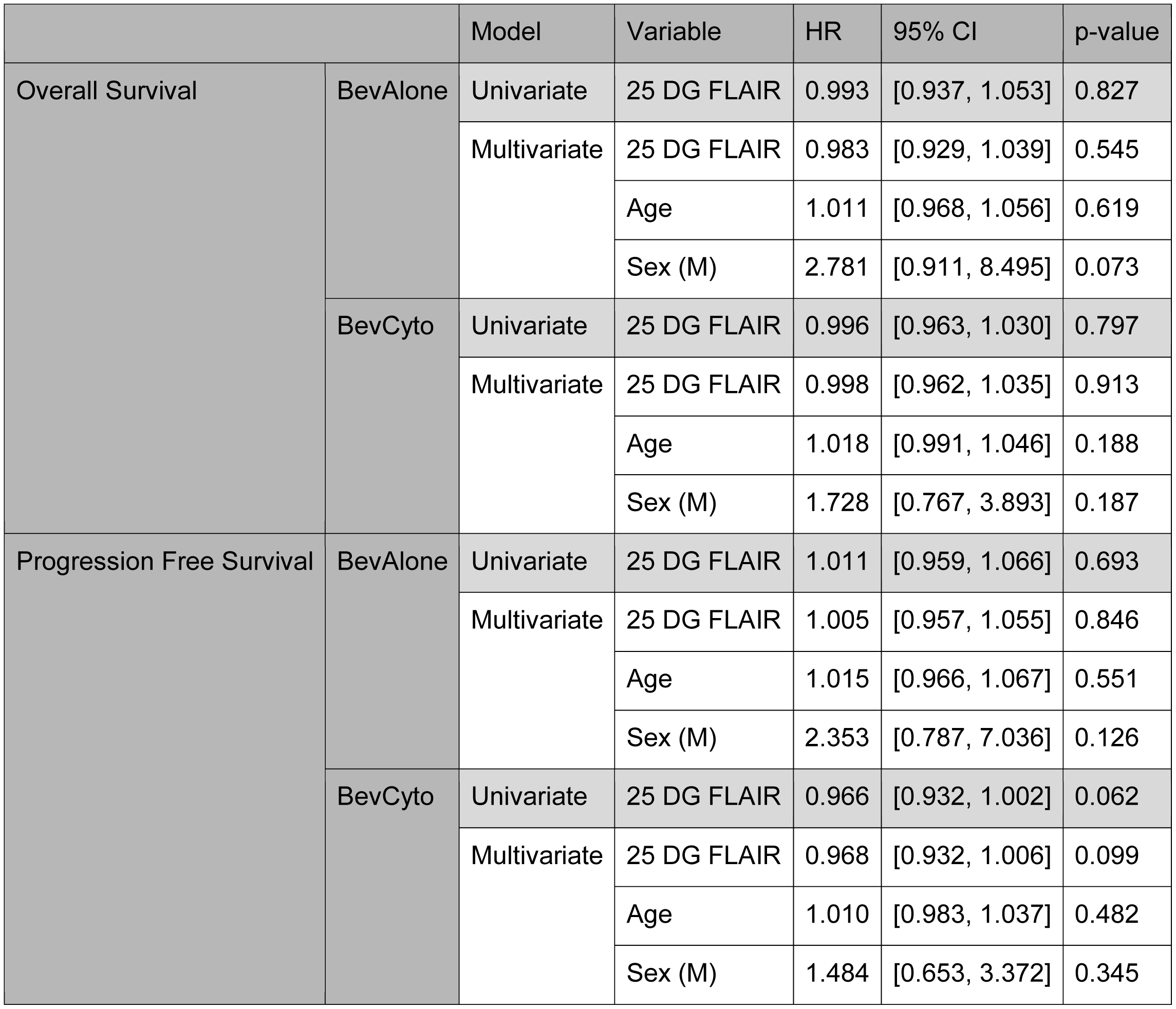
Cox proportional hazards regression analysis of overall survival and progression free survival using continuous DG_FLAIR_ scores, patient age at start of treatment, and patient sex. No significant p-values were observed.

